# Sequencing 4.3 million mutations in wheat promoters to understand and modify gene expression

**DOI:** 10.1101/2023.07.21.550110

**Authors:** Junli Zhang, Hongchun Xiong, Germán F. Burguener, Hans Vasquez-Gross, Qiujie Liu, Juan M. Debernardi, Alina Akhunova, Kim Campbell-Garland, Shahryar F. Kianian, Gina Brown-Guedira, Curtis Pozniak, Justin D. Faris, Eduard Akhunov, Jorge Dubcovsky

## Abstract

Wheat is an important contributor to global food security, and further improvements are required to feed a growing human population. New functional genetics and genomics tools can help us to understand the function of different genes and to engineer beneficial changes. In this study, we used a promoter capture assay to sequence 2-kb regions upstream of all high-confidence annotated genes from 1,513 mutagenized plants from the tetraploid wheat variety Kronos. We identified 4.3 million induced mutations with an accuracy of 99.8%, resulting in a mutation density of 41.9 mutations per kb. We also remapped Kronos exome capture reads to Chinese Spring RefSeq v1.1, identified 4.7 million mutations, and predicted their effects on annotated genes. Using these predictions, we identified 59% more non-synonymous substitutions and 49% more truncation mutations than in the original study. To show the biological value of the new promoter dataset, we selected two mutations within the promoter of the *VRN-A1* vernalization gene. Both mutations, located within transcription factor binding sites, significantly altered *VRN-A1* expression, and one reduced the number of spikelets per spike. These publicly available sequenced mutant datasets provide rapid and inexpensive access to induced variation in the promoters and coding regions of most wheat genes. These mutations can be used to understand and modulate gene expression and phenotypes for both basic and commercial applications, where limited governmental regulations can facilitate deployment. These mutant collections, together with gene editing, provide valuable tools to accelerate functional genetic studies in this economically important crop.

**Significance Statement:** We sequenced 4.3 million induced mutations in the promoters and 4.7 million in the coding regions of most wheat genes. We also show how this public resource can be used to understand gene function, modulate gene expression, and generate changes in valuable wheat agronomic traits.

## Introduction

Wheat provides one fifth of the calories and proteins consumed worldwide (1), and continuous improvements in wheat productivity and nutritional value are required to feed a growing human population. Crop improvement relies on natural or induced variation that alters the function or expression of genes controlling relevant traits. To induce mutations with detectable effects, wheat researchers used ionizing radiation in the late 1920s (2, 3) and expanded into chemical mutagenesis in the 1960s utilizing reagents such as ethyl methane sulfonate (EMS). EMS induces point mutations (G to A and the reciprocal C to T) that are less deleterious than the large deletions and chromosome breaks generated by radiation (4).

In early mutagenesis studies, researchers found that the use of induced mutations in polyploid wheat was not as effective in generating phenotypic changes as in diploid grasses, such as barley or rice. These researchers hypothesized that gene redundancy, resulting from polyploidy likely masks the cause of the phenotypic effects of the mutations (2), a hypothesis that was confirmed in more recent studies (5, 6). Duplicated genes (homeologs) result in reduced selection pressure and in new opportunities for sub-functionalization or elimination of duplicated genes. Differences between homeologs are limited in wheat due to the recent origin of the polyploid species. This results in higher levels of functional overlap among duplicated genes and reduced effects of induced mutations than in older polyploid species (6). Tetraploid wheat (2n = 28, genomes AABB) originated less than 0.8 MYA (7, 8), whereas hexaploid wheat (2n = 42, genomes AABBDD) originated ∼10,000 years ago (9, 10), providing little time for functional differentiation of homeologs.

The extensive functional overlap among homeologs allows the polyploid wheat species to tolerate higher doses of mutagen and more mutations per plant, which reduces the number of mutagenized plants required to saturate the gene space with mutations relative to diploid or old polyploid species (11). This property was used to develop saturated EMS mutagenized populations for both tetraploid and hexaploid wheat (5, 11–13). These populations were initially screened using DNA pools and the endonuclease *Cel*I, a technology called TILLING (targeting induced local lesions in genomes) (14). The *Cel*I assays were later replaced by high-throughput sequencing methods that can screen multiple genes simultaneously (15), and more recently by sequencing the coding regions of most wheat genes across the complete mutant population using exome capture (5). The latter study generated databases of 4.15 million sequenced EMS mutations in the tetraploid wheat cultivar Kronos and 6.42 million mutations in the hexaploid wheat cultivar Cadenza, which can be rapidly screened using web-based BLASTN searches (5).

These databases have been used to identify loss-of-function mutants in numerous wheat genes affecting heading time (16–21) and spike and grain development (22–31). However, loss-of-function mutations in coding regions can sometimes have negative phenotypic effects, limiting their use in wheat improvement. For these genes, it would be desirable to identify *cis*-regulatory variants affecting the level of expression or the timing or tissues where the gene is expressed, to generate more subtle phenotypic changes and reduced pleiotropic effects (32, 33).

Multiple examples of natural *cis*-regulatory single nucleotide polymorphisms (SNPs) affecting gene expression and valuable agronomic traits have been reported in plants. A single SNP in the *SH1* promoter affects its expression at the spikelet abscission layer and results in the non-shattering characteristic of domesticated rice (34). In the same species, natural SNPs in the promoters of genes *TGW2* and *GS5* affect gene expression and result in changes in grain size (35, 36). In wheat, a single SNP in the promoter of the *VRN-D1* gene has been associated with differences in heading time (37). In addition to natural variants, induced *cis*-regulatory mutations can be also useful for crop improvement (38). A collection of CRISPR edited alleles of the *CLV3* promoter in tomato has been shown to provide a continuum of fruit sizes (39), and edited weak promoters of the *CLE* genes in maize have been used to engineer quantitative variation in yield-related traits (40).

In this study, we sequenced the regulatory regions upstream of all the high-confidence annotated wheat genes (CS RefSeq v1.1, (41)) in a population of 1,513 EMS-mutagenized lines of tetraploid wheat variety Kronos, and identified 4.3 million induced mutations (99.8% accuracy). We also remapped the previous Kronos exome capture reads to the improved RefSeq v1.1 reference genome and predicted the effect of mutations in the annotated genes, significantly expanding the number of identified non-synonymous and truncation mutations. To show the value of this new tool, we selected two mutations in predicted regulatory regions of the *VRN-A1* gene and showed their significant effect on gene expression and number of spikelets per spike.

## Results

### Identification of EMS induced mutations in promoter regions of tetraploid wheat

We performed promoter captures for 1,535 Kronos EMS mutant lines using a previously published NimbleGen promoter capture for hexaploid wheat (42) and obtained an average of 135.3 M 150 PE reads per line (67.65 M read-pairs or approximately 20 Gb per line, *SI Appendix*, Method S1). After trimming, mapping to the wheat genome (RefSeq v1.1), and eliminating duplicated reads, we ran the MAPS pipeline (5) to identify mutations at four stringency levels (*SI Appendix*, Table S1 and Method S2).

These stringency levels differ by the minimum coverage (MC) required to call a heterozygous (HetMC) or homozygous (HomMC) mutation. At the lowest stringency, HetMC3/HomMC2 (at least 3 reads with the mutation in heterozygous and 2 reads in homozygous lines), we identified 4,657,354 mutations, 96.8% of which were G to A or C to T (henceforth, G>A/C>T or EMS mutations). At the HetMC5/HomMC3 higher stringency level, the number of called mutations decreased to 3,784,020 and the percent of EMS mutations increased to 99.1% (*SI Appendix*, Table S1), indicating a reduction in the error rate. Fig. 1 shows the number and type of mutations in the promoter regions for 15 randomly selected lines compared with the non-mutagenized Kronos control (at HetMC5/HomMC3).

**Figure 1.**
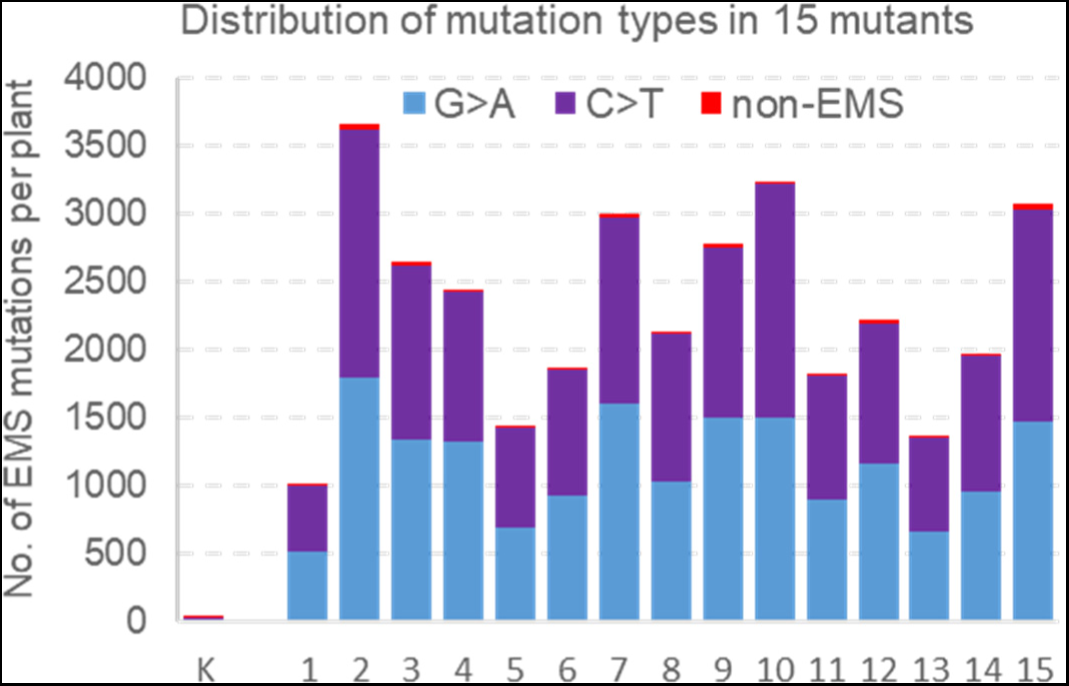
SNPs detected in the Kronos promoter capture. Mutations were called at HetMC5/HomMC3 for 15 random mutagenized Kronos lines and the non-mutagenized Kronos control (K). Blue and violet bars indicate EMS-type mutations (G>A/C>T) and red bars indicate non-EMS mutations. We identified 29 SNPs in the non-mutagenized Kronos control, which are likely sequencing errors or residual heterogeneity that our bioinformatics pipeline failed to identify and eliminate.

The percent of non-EMS mutations provides an estimate of the error rate, but it is not its final measure because non-EMS mutations are eliminated from the searchable databases resulting in a smaller error. However, G>A/C>T errors cannot be eliminated because they cannot be distinguished from real EMS mutations. To estimate this source of errors, we used the frequency of the EMS reciprocal transitions A to G or T to C SNPs (henceforth, A>G/T>C or non-EMS transitions), which have a similar frequency to the G>A/C>T SNPs among random mutations (5). The average predicted error rate calculated using this method was 0.15% at HetMC5/HomMC3, which is similar to the estimated errors at the same stringency level in the previously published study (0.18%) (5).

At this stringency level, we observed highly significant (*P* < 0.0001) correlations between the promoter and exome captures for the number of mutations (*R* = 0.854) across the 1,492 Kronos mutant lines shared between the two studies (*SI Appendix*, Table S2). Since these mutations were obtained from the same next-generation sequencing libraries, the significant correlation indicates that a large proportion of the variation among lines in number of mutations (*R^2^* = 0.73) is explained by either variation in the number of mutations present in each line (e.g. different exposure or penetration of EMS into seeds) and/or by variation in the quality of the sequencing libraries. The particular effect of the variation in sequencing library quality was more specifically reflected by the smaller, but still highly significant correlation between the two captures for the percentage of EMS (*R =* 0.154) and non-EMS transitions (*R* = 0.183, *SI Appendix*, Table S2).

### Determination of a new error threshold adjusted by sequencing library

The variability in %-EMS among lines suggested that our previous strategy of using the same error threshold for all the lines was not the best strategy to maximize the number of detected mutations while minimizing the error rate. Therefore, we developed a new strategy that adjusts the stringency level used to call mutations based on the quality of the library, estimated by the percent of non-EMS mutations. We found that a %-EMS ≥98% was sufficient to reduce the % of non-EMS transitions below 0.7 % even in the lowest quality libraries, so we called mutations in each library at the lowest stringency level that resulted in ≥98%-EMS (henceforth EMS98%, *SI Appendix*, Method S3).

A total of 940 among the 1,535 sequenced lines exceeded the EMS98% threshold at the lowest stringency level (HetMC3/HomMC2), yielding 3,166,422 EMS mutations (74% of the total EMS mutations). This result indicated that most of the libraries were of good quality. We selected 834,662 EMS mutations at HetMC4/HomMC3, 205,755 EMS mutations at HetMC5/HomMC3 and 80,522 EMS mutations at HetMC6/HomMC4. Finally, we eliminated 22 libraries that did not reach the EMS98% threshold even at the highest stringency level (HetMC6/HomMC4), resulting in 1,513 lines used in the promoter capture dataset.

By eliminating more SNPs from the low-quality libraries and selecting more from the good quality libraries, we were able to call 561,605 more EMS-type mutations than by using the HetMC5/HomMC3 across all libraries, while maintaining a very low estimated error (0.21%, *SI Appendix*, Table S1). The use of the new EMS98% method adjusted by library also reduced the correlation between the promoter and exome capture studies for percent EMS (from *R =* 0.154 to *R =* 0.034) and non-EMS transitions (from *R =* 0.183 to *R =* 0.119), suggesting a reduced effect of the differences in sequencing library quality on the mutation discovery process (*SI Appendix*, Table S2).

In summary, using the new EMS98% method, we identified 4,287,361 EMS-type promoter mutations from 1,513 libraries (2,834 EMS mutations per line) with an estimated error rate of 0.21% (Table 1). Based on an estimated mapping space of 102,378,005 bp (quality ≥ 20, coverage ≥ 3), we estimated a mutation density of 41.9 EMS mutations/kb across the complete population, or 23.8 EMS mutations per Mb per individual line (Table 1).

**Table 1.** Summary of promoter and exome capture mutations identified using the EMS98% method. The comparison between parameters for the promoter capture obtained by methods EMS98% adjusted by library and HetMC5/HomMC3 global adjustment is provided in *SI Appendix*, Table S1.

| Trait | Promoter | Exome |
| --- | --- | --- |
| Number of lines used (overlap 1,465) | 1,513 | 1,521 |
| Total uniquely mapped SNPs | 4,345,625 | 4,748,394 |
| **Uniquely mapped EMS-type mutations \*** | **4,287,361** | **4,690,454** |
| Valid mapping space (quality $\geq 20$, min. coverage 3, no RH) | 102,378,005 | 131,190,164 |
| Avg. EMS-type mutations per kb (population) | 41.9 | 35.8 |
| Avg. EMS-type mutations/line * | 2,833.7 | 3,083.8 |
| % EMS-type * | 98.7 % | 98.8 % |
| % Heterozygous for EMS SNPs (expected 66.66% at M<sub>2</sub>) * | 64.67 % | 65.07 % |
| % Heterozygous for non-EMS SNPs | 99.99 % | 99.99 % |
| Non-EMS-type transitions (estimated error) *† | 0.21 % | 0.34 % |
| Residual heterogeneity (RH) SNPs | 40,823 | 73,239 |
| % RH SNPs | 0.93 % | 1.52 % |
| % Heterozygous in RH | 14.7 % | 26.8 % |
| % EMS-type in RH | 16.9 % | 23.0 % |
| Non-EMS-type transitions in RH | 25.4 % | 29.9 % |

\* Excluding residual heterogeneity (RH) and deletions.
† Estimated from the percent of reciprocal A>G and T>C transitions among total mutations.

### Remapping exome capture to RefSeq v1.1 using the EMS98% error method

The previous exome capture data was mapped to a fragmented wheat genome with limited annotation (5). Since there are no conversion tables from previous assemblies to RefSeqv1.0, we decided to remap the reads to the new CS RefSeq v1.1 using the new error threshold, and to re-annotate the mutation effects using the new gene models. This involved the remapping of the exome captures of 1,535 Kronos lines and running the MAPS pipeline. We called SNPs using the same four stringency levels as in the promoter capture described above, and then used the new EMS98% threshold method to select high-confidence mutations.

The use of the EMS98% threshold resulted in the elimination of 14 libraries that showed less than 98%-EMS even at the highest stringency level (HetMC6/HomMC4), but still yielded 537,629 more EMS mutations relative to the previous HetMC5/HomMC3 threshold across all libraries (*SI Appendix*, Table S1). Using the new threshold, we identified 4,690,454 unique EMS-type mutations from 1,521 libraries with an estimated error rate of 0.34 % (Table 1). This represents an average of 3,084 EMS mutations per line, and a mutation density of 36.7 EMS mutations/kb across the complete population (Table 1, exome capture).

We determined mutation effects using the Variant Effect Predictor (VEP) program (43) (*SI Appendix*, Method S4). VEP was also used to predict Sorting Intolerant from Tolerant (SIFT) scores (44), which predict the potential impact of amino acid substitutions on protein function. The improved effect prediction tools, together with the improved gene models of CS RefSeq v1.1 increased the number of annotated missense variants to 1,637,961 (59 % increase) and of truncation mutations to 113,880 (49 % increase) relative to the previously published study (5) (*SI Appendix*, Table S3). As a result of the additional annotated effects, the number of gene models with at least one missense mutation increased 22.9 %, and the number of genes with predicted truncations (premature stop codons plus splice site mutations) increased by 34.9 %, relative to previous results (5) (*SI Appendix*, Table S3).

Some of the mapped reads in the exome capture extend upstream of the start codon, so we expected some overlap between the mutations detected in the promoter and exome capture studies. We used this overlap to confirm the correct identity of the lines in both captures. We identified 174,677 mutations in 1,465 lines shared between the promoter capture and exome capture (*SI Appendix*, Table S4), confirming the correct tracking of the sequencing libraries and file identification numbers in the two experiments, and the quality of the called mutations.

### Distribution of EMS mutations across and along the wheat chromosomes

The number of EMS mutations per chromosome was highly correlated with the number of annotated genes for both the exome (*R* = 0.942) and promoter captures (*R* = 0.926, *SI Appendix*, Table S5). These results indicate that the probes captured mainly the gene regions as intended, and that the differences in the number of SNPs across chromosomes were mainly driven by their differences in gene content.

The EMS mutation density is relatively uniform, so the average number of detected mutations per kilobase is directly correlated with the length of DNA captured for a particular region. Since the promoter and exome captures are focused on genes, and wheat genes are more abundant in the distal regions of the chromosomes, we expected more mutations in those regions. To visualize the distribution of EMS mutations within chromosomes, we generated a circle graph including both the promoter and remapped exome capture data (Fig. 2). This graph showed that the EMS mutations in both studies are, as expected, more abundant towards the distal regions of the chromosomes, reflecting the higher gene density in the distal regions of the wheat chromosomes (45).

**Figure 2.**
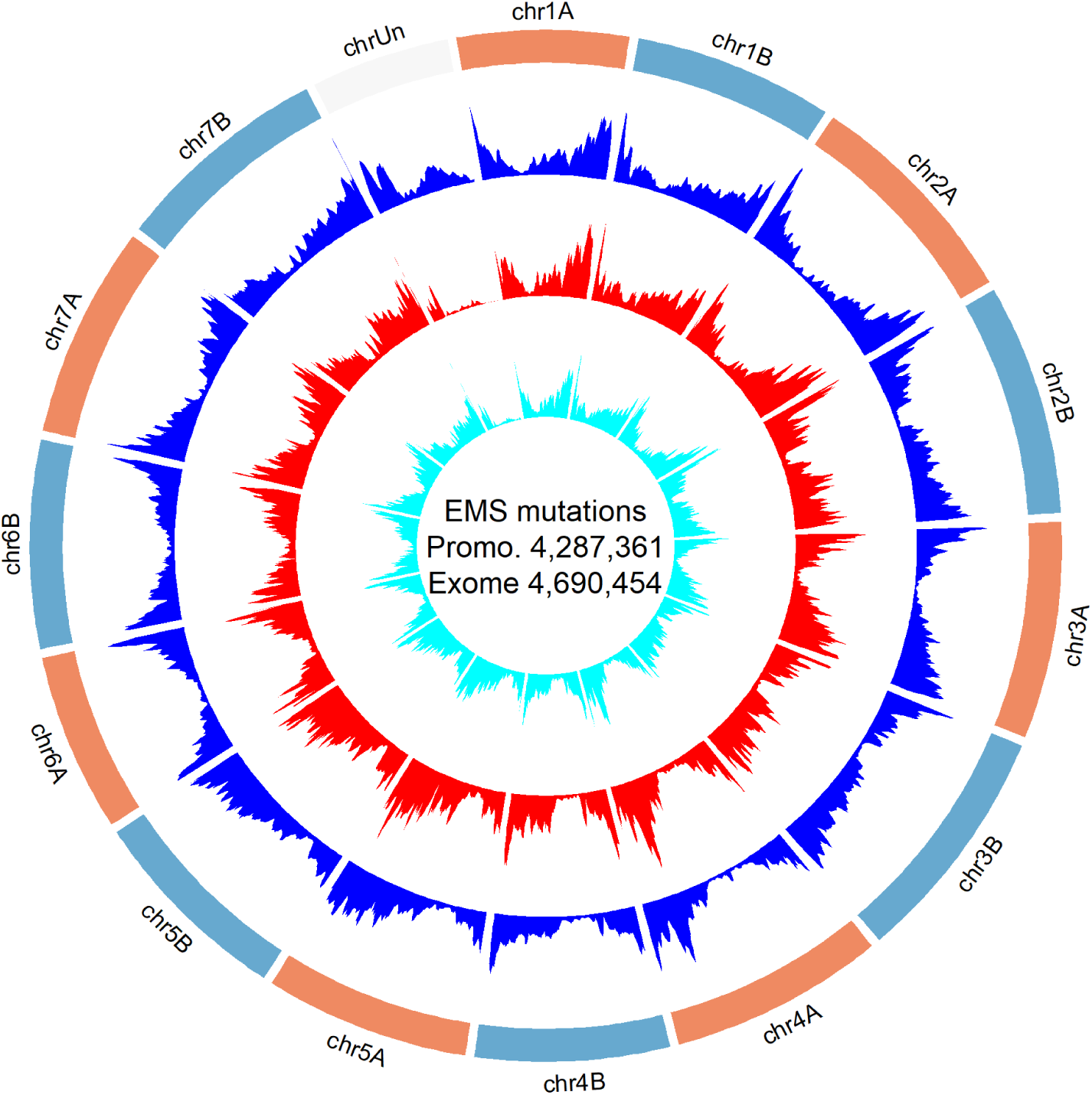
Genome-wide positions of identified mutations in the promoter and exome capture. The blue track indicates gene density (high confidence gene from RefSeq v1.1 annotation), the red track indicates EMS mutation density in the promoter capture, and the cyan track indicates the EMS mutation density in the reanalyzed exome capture.

### Residual heterogeneity (RH) regions

The previous Kronos exome capture revealed that 1.64 % of the detected SNPs were not induced by EMS mutagenesis but were instead originated from residual heterogeneity present in the mutagenized Kronos seed (5). We identified these regions by their higher proportion of linked non-EMS mutations, higher SNP density, higher proportion of homozygous SNPs, and by the presence of the same SNPs in multiple individuals. We combined those four criteria in one index and used a bioinformatics pipeline developed in a previous study (5) to identify and eliminate mutations in the predicted RH regions. This filter resulted in the elimination of 40,823 SNPs in the promoter capture (0.93 %) and 73,239 SNPs in the exome capture (1.52 %, Table 1). These percentages of RH are consistent with pooled seeds from different plants after six generations of self-pollination, a common practice in wheat breeding.

### Percent of heterozygous SNPs

We collected the DNAs used in this study from M_2_ plants, so we expected 1/4 of the plants to be homozygous for the wild-type allele, 1/4 homozygous for the mutant allele, and 1/2 heterozygous. Since we can only identify the mutant alleles, we expect 2/3 of the identified mutations to be heterozygous (66.7 %). However, we initially observed 68.1 % heterozygous EMS mutations in the promoter capture and 68.2 % in the reanalyzed exome capture. This excess heterozygosity is likely caused by the low threshold used by the MAPS program to classify mutations as heterozygous (5). Even when a single wild-type read is mapped to a mutation site, MAPS calls that mutation heterozygous even if it is at a very low frequency relative to the reads showing the mutation. This can happen, for example, when a read from a closely related homeolog is erroneously mapped to the region carrying the mutation and is not eliminated by the MAPS pipeline.

To correct these errors, we used a heterozygous filter pipeline developed in the initial exome capture project (5). This filter reclassifies heterozygous mutations as homozygous when the frequency of the wild-type allele is less than 15 % of the reads (5). After we applied this filter, the average percentage of heterozygous EMS mutations (%-het) was reduced to 64.7 % for the promoter capture mutations and to 65.1 % for the exome capture mutations.

Unexpectedly, 100 lines showed an average proportion of heterozygous mutation (>90 %) in both the promoter and exome captures that was much higher than expected for M_2_ plants (66 %, Fig. 3A-B). These 100 lines showed similar average % EMS (98.4 – 98.7 %) as the other 1,365 shared lines (98.6 – 98.7 %), but they had on average 17 % more mutations than the rest of the lines both in the promoter and exome captures (*P <* 0.0001, Fig. 3C). We hypothesize that the M_2_ plants with >90 % heterozygous mutations originated by hybridization between two mutant M_1_ gametes. The maximum proportion of shared mutations between each of the 100 lines with >90 % heterozygous and any of the other 1,365 lines was 0.51 % indicating that none of the 100 lines originated from crosses with any of the other sequenced Kronos mutant lines (which will be expected to share ∼50% of the mutations). Therefore, we hypothesize that hybridization occurred between gametes from chimeric mutant tillers from the same M_1_ plant or between mutant plants not included in our subset of sequenced lines. The percent cross-pollination in wheat is usually small (46), but the increased sterility of the mutant lines likely increased this probability.

**Figure 3.**
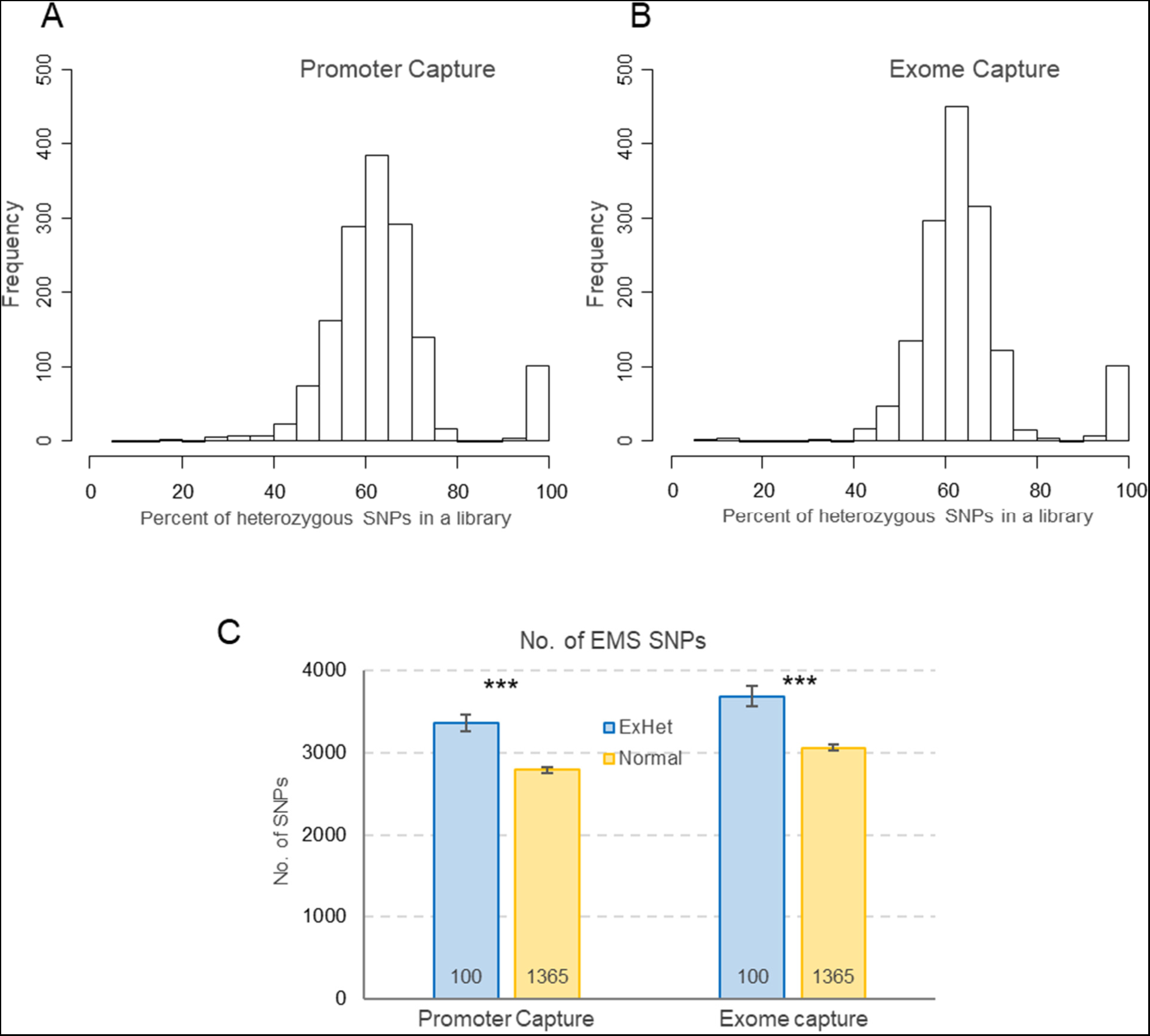
Characterization of lines with higher-than expected levels of heterozygous EMS mutations. Distribution of the proportion of heterozygous mutations per line in **a** promoter and **b** exome capture populations. **c** Comparison of the average number of SNPs between 100 lines with >90 % heterozygous mutations (ExHet, excess heterozygous) and 1,365 lines with ≤90 % heterozygous among the 1,465 lines shared between the promoter and exome capture datasets. *P* values are from two-sided Kruskal-Wallis tests and error bars are s.e.m. *** = *P* < 0.001.

### EMS “preferred” G residues and level of mutagenesis saturation

We analyzed the 10 bp flanking the mutated G sites in the promoter capture to determine if there was a preferred sequence context for the mutations. We found a relatively high frequency of C bases at position +1 downstream of the mutated G, and of G at position –1 and +2 relative to the mutated G; whereas a negative bias for T was observed at position –1 (Fig. 4A). These preferences are similar to those reported in the previous exome capture study (5), and indicate that not all the G bases in the genome have a similar probability of being mutated by EMS.

**Figure 4.**
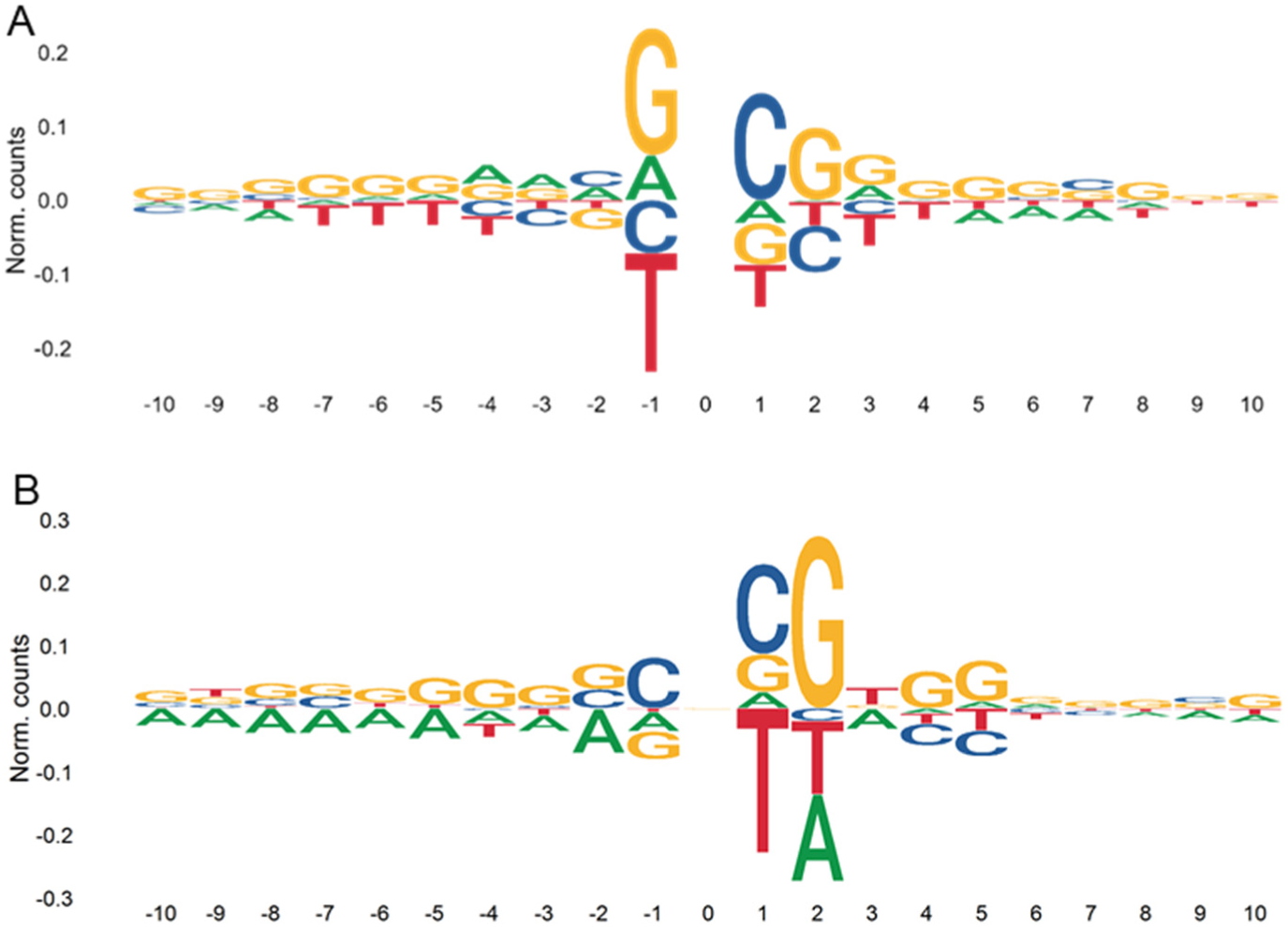
Sequence context of G>A EMS mutations and C>A errors. **a**. Sequence preference in regions flanking EMS-type G>A calculated based on all EMS mutations **b**. C>A sequence context was calculated based on the 60 sequencing libraries in both captures with the highest %C>A/G>T. The x-axis indicates the number of nucleotides upstream (negative) and downstream (positive) from the mutated site.

Although this variation is likely continuous, we artificially divided them into two hypothetical populations of EMS-accessible and EMS-inaccessible GC bases to simulate a Poisson distribution and estimate the average proportion of accessible GC positions. We wanted to use this value to calculate the percent of the EMS-accessible GC positions for which we already have mutations in our population, a parameter we will refer hereafter as the percent saturation. This parameter is useful to decide if it is worth sequencing additional wheat mutant lines treated with EMS or if it is better to switch to a different mutagen that can access different sequences in the genome. As the percent saturation increases, additional sequencing results in diminishing returns because the proportion of mutations found in more than one line (henceforth, duplicated mutations) increases.

Among the large number of mutations identified in this study, we found 291,577 EMS mutations in the promoter capture (PC) and 281,686 in the exome capture (EC) that were present in two lines. Those numbers decayed rapidly for mutation shared by three (PC= 31,539 and EC= 25,214) or four (PC= 4,503 and EC= 3,228) lines (*SI Appendix*, Table S6), following an approximate Poisson distribution. To estimate the proportion of EMS accessible GCs, we first tested different means (λ = average mutations per site across the population) to identify the Poisson distribution that better fit the observed data. We found that λ= 0.182 and λ= 0.153 minimized the differences between observed and expected values for the promoter and exome captures, respectively (*SI Appendix*, Table S6 and Method S5).

Using λ = 0.182, we estimated the existence of 23,556,929 “accessible” GC sites among the 47.9 M predicted GC in the total mapping space of the promoter capture (102.4 Mb, GC content of 46.8 %). Using a similar procedure, we estimated that, on average, 50.7 % of the GC sites in the exome capture space were accessible to EMS (*SI Appendix*, Table S6). The lower estimate for this parameter presented in our previous exome capture paper (5) was due to a calculation error, and an “erratum” has been submitted showing similar values to those presented here. In summary, these estimations suggest that approximately half of the GC sites in the sequenced space were accessible to the EMS treatment performed in this study.

We then used the estimated EMS-accessible GC to calculate the %-saturation values (mutated sites / accessible GC sites). Only 14.2 % of the EMS-accessible GC in the exome capture and 16.6 % in the promoter capture are covered by EMS mutations in at least one of the sequenced lines in the mutant population. The larger value observed in the promoter capture reflects the higher mutation density in the promoter capture (41.9 mutations / kb) than in the exome capture (37.7 mutations per / kb). The percent-saturation values estimate the probability that a new mutation will be the same as a previously identified one, a probability that can be also estimated by dividing the number of duplicated mutations by the total number of EMS mutations. These two independent estimates are almost identical validating the Poisson simulation (*SI Appendix*, Table S6).

### Functional characterization of selected mutations

To show the value of the sequenced promoter mutations, we explored the 2-kb upstream of the vernalization gene *VRN1*, which is a central regulator of heading time (47–49) and spike development (24) in wheat. We first delimited conserved regions in the promoter by aligning *VRN1* orthologs from wheat, barley, rice, maize, sorghum, and *Brachypodium* using T-coffee (https://tcoffee.crg.eu/). We focused on two conserved regions (Figs. 5A and E) including predicted binding sites for transcription factors SQUAMOSA PROMOTER BINDING PROTEIN LIKE (SPL) and LFY (Fig. 5B and F, respectively).

**Figure 5.**
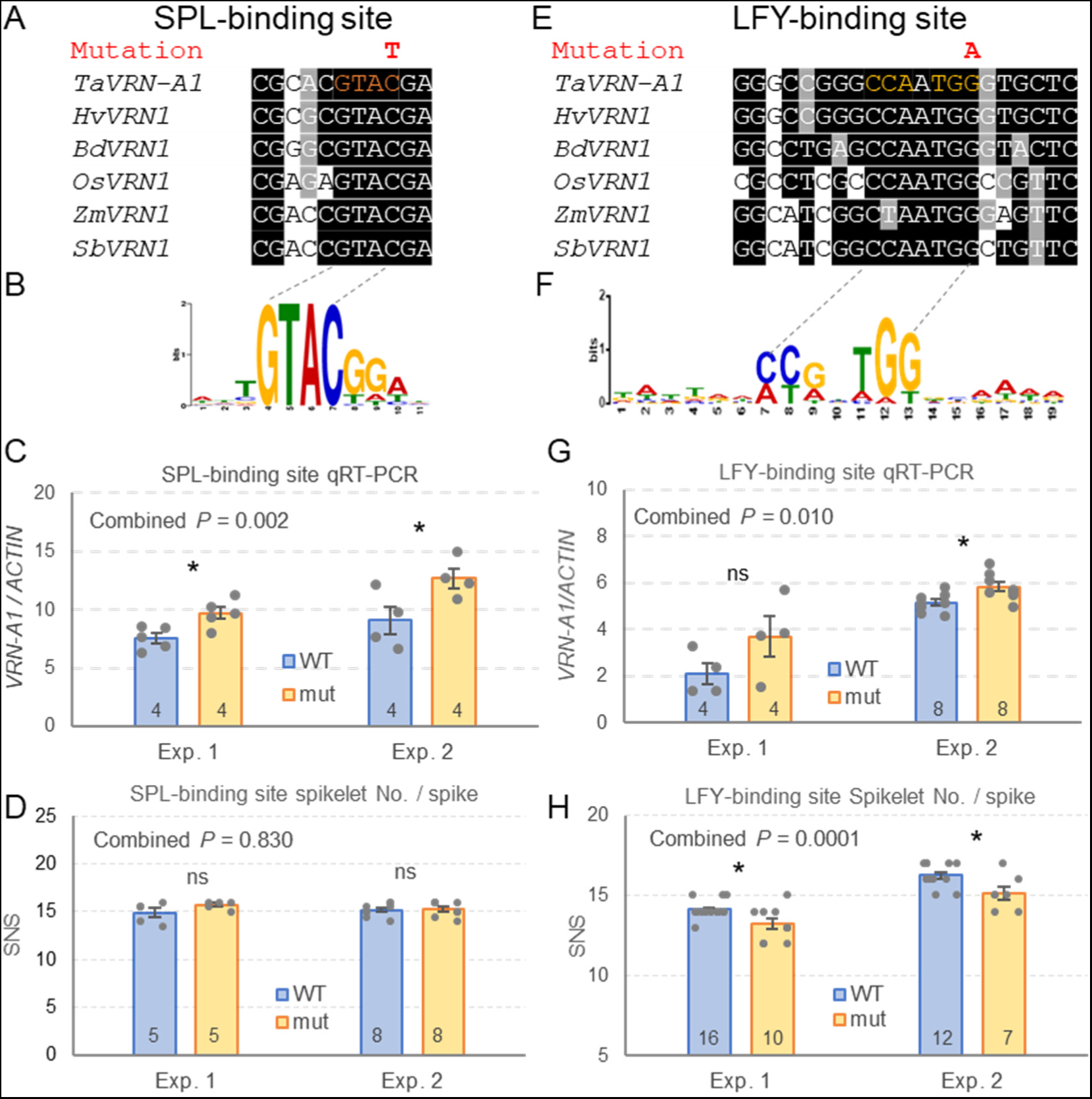
Effect of promoter mutations on *VRN-A1* expression and spikelet number per spike (SNS). **a-d** Mutation in a conserved putative SPL-binding site. **e-f** Mutation in a conserved putative LFY-binding site. **a** and **e** Conserved regions in the *VRN1* promoter in wheat (*Ta*), barley (*Hv*), *Brachypodium distachyon* (*Bd*), rice (*Os*), maize (*Zm*), and sorghum (*Sb*). **b** and **f** Sequence preference logo for SPL and LFY transcription factors. **c** and **g** Effect of mutations in *VRN-A1* expression. **d** and **h** Effect of mutations on the number of spikelets per spike (SNS). ns= not significant, * = *P* < 0.05, ** = *P* < 0.01, *** = *P* < 0.001. Numbers at the bottom of the bars indicate the number of biological replicates (grey dots) and error bars are s.e.m. ANOVAs combining the two experiments are available in *SI Appendix*, Tables S7 to S10, and the combined *P* value for genotype is indicated in the top left corner of each graph.

We first selected a C to T mutation in Kronos mutant K4679 located within the predicted SPL core binding site (GTAC) (50). This mutation, located –368 bp upstream from the *VRN-A1* start codon, changes the core binding motif GTA<u>C</u> to GTA<u>T</u> in a position that is well conserved across grass species (Fig. 5A). Since SPL proteins are known to affect panicle development in rice (51–53), we explored the effect of the selected mutation on the expression of *VRN-A1* in young developing spikes (5 mm long) dissected from tetraploid wheat Kronos plants. The expression of *VRN-A1* in the mutant line across two experiments was on average 33.6 % higher than in the wild-type sister line (*P* = 0.002, Fig. 5C, *SI Appendix*, Table S7), confirming that this mutation affects *VRN-A1* expression. However, we did not detect significant differences in spikelet number per spike (SNS, Fig. 5D) or heading time (*SI Appendix*, Table S8) in this particular mutant line.

We also found a mutation in Kronos mutant K2944 within a predicted LFY binding site (54) located between positions –285 to –281 upstream from the *VRN-A1* start codon. This mutation resulted in a change from the conserved CC(A/G)n(T/A)G<u>G</u> binding site (54) to CC(A/G)n(T/A)G<u>A</u>, in a position that was conserved across the analyzed grass species (Fig. 5E and F). We then studied *VRN-A1* transcript levels at the early stages of spike development (lemma primordium present, Waddington scale W3.25, ∼2 mm long) by qRT-PCR in two independent experiments (Fig. 5G, *SI Appendix*, Table S9). The differences in expression were significant only in the second experiment with a larger number of replications (Fig. 5G), but the trend was the same in both experiments resulting in a significant difference in the combined ANOVA using experiments as blocks (*P* = 0.010, *SI Appendix*, Table S9). On average, we observed a 27% higher *VRN-A1* expression in the line with the mutation in the LFY binding site than in the wildtype. This change was associated with an average reduction of 6.6 % in the number of spikelets per spike relative to the sister lines with the wild-type allele (*P* = 0.0001, Fig. 5H, *SI Appendix*, Table S10). No significant differences were observed for heading time and leaf number (*SI Appendix*, Table S10).

In summary, these examples suggest that the catalogue of sequenced promoter mutations developed in this study will be useful for regulatory variants to manipulate the levels of gene expression and to fine tune phenotypic changes in wheat.

## Discussion

The large and repetitive nature of the polyploid wheat genomes delayed the initial genome sequencing efforts relative to other diploid crop species with smaller genomes. However, since the release of the first hexaploid wheat reference genome (41), multiple complete wheat genomes have been sequenced (55), exome sequencing data has been completed for ∼1000 wheat accessions (56), and expression data has been generated from multiple tissues and germplasm (57, 58). These public genomics resources, together with more efficient wheat transformation technologies (59, 60), have significantly expanded the opportunities for wheat research and improvement to more groups around the world. In this study, we report the development of a public database of 4.3 million sequenced mutations in the promoters of most genes in durum wheat, and the expansion and improved annotation of the sequenced mutations in the coding gene region from the same species.

## Expanded exome capture mutant dataset with improved annotations

The first exome capture sequencing studies focused on mutant populations were developed for tetraploid cv. Kronos (1,535 lines) and hexaploid cv. Cadenza (1,200 lines) in 2017 (5), before the release of the first wheat genome reference CS RefSeq v1.0 (41). In those studies, reads were aligned to fragmented scaffolds assembled using Illumina short reads generated from flow-sorted chromosome arms (Chromosome Survey Sequencing [CSS]) (61). In spite of these initial limitations, the two mutant wheat collections have been extensively used by the international wheat research and breeding community, with >20,000 mutant seed samples distributed until March 2023 by the University of California (USA) and the John Innes Centre (UK) (plus an unknown number from five other locations that received complete Kronos mutant populations, including laboratories in Australia, Canada and China (5)).

To increase the value of these resources, we remapped the reads from the two exome captures to the CS RefSeq v1.0 released in 2018 (41), and predicted the effects of the mutations using the improved gene annotation from CS RefSeq v1.1. Cadenza reads from the 1,200 lines were called using the DRAGEN system (62), whereas Kronos reads from the 1,535 lines were remapped using the MAPS pipeline (15) as part of this study. Both datasets are available in the GrainGenes Genome browser (https://wheat.pw.usda.gov/GG3/genome_browser) for CS RefSeq v1.0. The Cadenza mutants are also available in ENSEMBL, whereas the expanded Kronos mutations described in this study will be incorporated in the next ENSEMBL release 105 (VCF files have been submitted).

The remapped Kronos exome capture detected 561,605 additional mutations compared with the original study (5), while maintaining a very low estimated error (0.34 %). This was achieved by using a stringency level adjusted by library rather than the uniform HetMC5/HomMC3 used in the previous study. This change was motivated by the discovery of a significant correlation in %-EMS between the exome and promoter captures in Kronos, which were generated using the same sequencing libraries. This significant correlation suggested that the %-EMS was significantly affected by the quality of the libraries. By adjusting the stringency level based on the quality of the library, we were able to eliminate more mutations from the poor-quality libraries (lower %-EMS) and to extract more mutations from the good quality ones, which greatly exceeded the problematic ones. This new method reduced the correlation between the promoter and exome capture studies for %-EMS (*SI Appendix*, Table S2), suggesting a reduced effect of the differences in sequencing library quality on the selected mutations.

The characterization of the non-EMS mutations (*SI Appendix*, Table S11), revealed that they were all highly heterozygous (>99.9 %). This result suggested that these were errors originated in the M_2_ plants or DNA, possibly during library construction or sequencing. We also observed an excess of C>A/G>T changes relative to the other non-EMS changes both in the promoter (47.9 %) and exome capture (36.8 %, *SI Appendix*, Table S11). One potential cause for a high proportion of C>A/G>T errors is DNA oxidation during the acoustic shearing step of sequencing library preparation, particularly when the DNA contains reactive contaminants from the extraction process (63). Once a guanine is converted to 8-oxoG via oxidation, it can pair with both cytosine and adenosine during PCR, leading to C>A/G>T transversions, that are more frequent in the C<u>C</u>G context (63). We found a similar context (GC)<u>C</u>G (Fig. 4B) in this study, using 60 lines with high C>A/G>T changes in both the promoter and exome captures. These results suggest that DNA oxidation during sonication may have contributed to the higher proportion of C>A/G>T changes relative to other errors, and to the correlation between the promoter and exome captures in the % EMS.

In summary, the detection of 561,605 additional high-confidence EMS mutations in the new exome capture, together with the improved gene annotation in CS RefSeq v1.1 relative to the previous Chromosome Survey Sequencing (5), resulted in a significant increase in the number of identified missense (59 %) and truncation (49 %) mutations, greatly increasing the value of this functional genetics resource.

### 4.3 million sequenced mutations in the promoter regions of wheat genes

The most significant contribution of this study is the database of 4.3 million mutations identified in the promoter regions of the annotated wheat genes. These promoter regions include *cis*-regulatory elements that control the location, growth stage, and levels of gene expression, and are frequently associated with open chromatin regions (64). Mutations in open chromatin regions, including promoter regions, have played an important role in plant and animal domestication (65–67), and explain a large amount of heritable phenotypic variance in diverse complex agronomic traits in maize and wheat (68, 69).

Changes in regulatory regions are frequently associated with more subtle phenotypic changes, and their pleiotropic effects are usually smaller than mutations in coding regions (32, 33). These properties of regulatory mutations are desirable for breeding objectives, where drastic phenotypic changes are frequently too disruptive for commercial use. The modulation of the expression of meristem regulatory genes *WUSCHEL* and *CLV3* to generate gradual variation in tomato fruit size (39) provides a good example of these potential applications.

One limitation for the use of EMS-induced mutations to modify gene expression, is that *cis*-regulatory elements are usually small, and a high mutation density is required to hit the small conserved core motifs. Even with the relatively high mutation density of the Kronos promoter capture (41 mutations/kb or one mutation every 25 bp), the probability of finding a mutation in a small 6 bp target remains low (∼25 %). Fortunately, the estimated level of EMS mutation saturation in our promoter capture database is relatively low (16.6 %), suggesting that mutation density can be further increased by sequencing additional EMS mutagenized individuals from this population. In addition, the development of sequenced promoter captures in other mutagenized wheat cultivars will also increase the chance to find a mutation within a specific *cis*-regulatory element.

The sequenced mutant database for wheat promoter regions generated in this study can be complemented by CRISPR, which produces targeted mutations, and can be used to edit binding sites not covered by the EMS induced mutations. CRISPR has also the advantage that efficient guide RNAs can be used to target simultaneously *cis*-regulatory elements in the different homeologs, and that off-target mutations are less frequent than in mutagenized plants. Genome editing is now extensively used to study gene function in plants and multiple examples of promoter editing in crop species have been reported recently (reviewed in (38)). However, sequenced EMS mutant databases also have some advantages, making these two approaches complementary. First, EMS mutations are not currently under government regulations in any country, facilitating their rapid incorporation into commercial cultivars. By contrast, genome editing is still facing different levels of regulation in different countries, which can be expensive and can be problematic for a globally traded crop such as wheat. Second, the generation of transgenic wheat plants still requires access to a sophisticated laboratory, which may be prohibitive for a developing country or a small wheat breeding program. However, as regulations for edited plants are relaxed, costs of transgenic plants are reduced, and more efficient promoter editing CRISPR technologies emerge (38), the use of genome editing to modify regulatory gene regions in commercial crop plants is expected to increase. Meanwhile, the wheat sequenced mutant databases presented in this study is a useful and inexpensive resource for rapid deployment of mutations in commercial wheat varieties.

A limitation for the use of promoter mutations generated by EMS or CRISPR in wheat, is that sometimes the effect of the mutation in one genome can be masked by the wild-type expression of the homeologous genes. In this case, the promoter mutant can be crossed with a loss-of-function mutation in the homeologous gene to magnify its phenotypic effects. In addition, a couple of backcrosses with the non-mutagenized Kronos may be required to reduce variability and increase the power to detect those phenotypic effects. Another limitation for both approaches is that the annotation of the promoters in most wheat genes is almost non-existent. Fortunately, ATAC-seq (64, 70), DAP-seq (71, 72) and CHIP-seq (21, 73) experiments in wheat are leading to improvements in the annotation of promoter and *cis*-regulatory element of some important genes. ATAC-seq data from wheat roots and leaf protoplasts (64, 70) have been incorporated into the public genome browser for CS RefSeq v1.0 in GrainGenes (https://graingenes.org/GG3/genome_browser) together with the promoter mutations reported here. Combining these tracks, the users can prioritize promoter mutations located in open chromatin regions.

## Characterization of mutations in the *VRN-A1* promoter

In this study, we explored the effect of induced mutations in conserved regions of the *VRN1* promoter on heading time and SNS as an example of the value of this sequenced mutant collection. *VRN1* encodes a MADS-box protein that plays a critical role in the regulation of the transition of the shoot apical meristem (SAM) to the reproductive phase (49, 74). The ancestral *VRN1* allele for winter growth habit requires long exposures to low temperature (vernalization) to be expressed, making *VRN1* a central gene in the flowering pathway in the temperate grasses (49, 74–77). However, mutations in the *VRN1* promoter or deletions in the first intron eliminate the vernalization requirement resulting in a spring growth habit (47, 48, 78, 79). The Kronos *VRN-A1* allele targeted in this study has a large intron deletion that eliminates the vernalization requirement.

Loss-of-function mutations in *VRN1* not only delay Kronos heading time but also increase the number of spikelets per spike (SNS), indicating a role in the regulations of the transition of the inflorescence meristem into a terminal spikelet (24). Plants with combined loss-of-function mutants in *VRN1* and its closest paralog *FUL2* cannot form spikelets and have an indeterminate spike, indicating an essential role of these two genes in the formation of both terminal and lateral spikelets (24). Overexpression of rice *MADS15* (the homolog of *FUL2*) reduces the number of primary branches in the rice panicle (80), suggesting a conserved role of these MADS-box genes in inflorescence development in grasses.

We focused on the effect of two EMS mutations located within conserved regions of the *VRN-A1* promoter, which potentially affect SPL and LFY predicted binding sites. In Arabidopsis, these two transcription factors are known to bind directly to the promoter of *AP1* (81, 82), which is a homolog of wheat *VRN1* (83). SPLs are plant-specific transcription factors that bind to a GTAC core sequence that is conserved from Arabidopsis (82) to wheat (84), and are known to affect flowering time (82) and inflorescence architecture (51, 85). The mutation in the conserved SPL-binding site in the *VRN1* promoter resulted in a small but significant effect on *VRN1* transcript levels in early spike development (Fig. 5C), which was not associated with significant changes in heading time or SNS (Fig. 5D). We are currently crossing the SPL-binding site mutant with a *VRN-B1* mutant, to test if the expression of the homeologous gene is masking the phenotypic effect of the mutations.

LFY plays an important role in the specification of the floral meristem in Arabidopsis, and a separate role in the transcriptional activation of *AP1* (81). LFY loss-of-function mutations in rice show increased transcript levels of *MADS14* and *MADS15* (rice homologs of *AP1* and *VRN1*) and reduced number of primary branches and spikelet numbers, suggesting that LFY acts as a transcriptional repressor of these MADS-box genes during early panicle development (86). Consistent with the results of studies in rice, the Kronos mutant for the LFY-binding site in the *VRN1* promoter was associated with a slight but significant upregulation of *VRN1* during the early stages of spikelet development (before terminal spikelet formation) (Fig. 5G) and a significant reduction in the number of spikelets per spike (Fig. 5H). This result indicates that LFY is involved in the regulation of the transition of the inflorescence meristem to a terminal spikelet.

Interestingly, the LFY-binding site mutant showed no significant differences in heading time or leaf number from the wild-type plants (*SI Appendix*, Table S10), which suggests that this mutation has limited effect on the timing of the transition of the SAM from the vegetative to reproductive stage or in the duration of the elongation phase. Taken together, these results indicate that mutations in the *VRN1* regulatory regions can be used to separate the pleiotropic effects of this gene on heading time and SNS.

New methods, such as DAP-seq (87) are generating massive amounts of predicted transcription factor (TF) binding motifs, which will require functional validation. Sequenced promoter captures of mutant populations, as the one presented here for tetraploid wheat, can provide a general and inexpensive resource to validate these predicted binding sites. The two mutations in the promoter of the wheat *VRN1* gene described in this study also exemplify the potential of this genomic tool to alter gene expression and modulate phenotypic effects.

The rapid increases in wheat production and nutritional value required to feed a growing human population rely on natural or induced genetic variants affecting useful agronomic traits. We provide here a database of 4.3 million sequenced induced mutations in the promoter regions of most wheat genes, which can be used to validate regulatory regions and modify gene expression and associated phenotypes. In addition, the expanded database of 4.7 million mutations in gene coding regions and their improved annotations provide a useful tool to study gene function and generate allelic variants not present in nature. These publicly available sequenced wheat mutant lines have the potential to accelerate and democratize functional genetic studies in a crop species that is critical for global food security.

## Materials and Methods

### Promoter capture and sequencing

Library preparation, capture and sequencing are described in detail in *SI Appendix*, Method S1. The wheat promoter capture (42) probes were ordered from Roche (SeqCap EZ Prime Developer Probes 96 Reaction, catalog number 8247633001). For the capture, 24 DNA libraries (125 ng/sample, 3 µg in total) were pooled and mixed with blocking oligos, denatured at 95 °C for 10 minutes and hybridized to the probes for 70 h at 48 °C.

The captured DNA was amplified for ten cycles using KAPA HiFi HotStart ReadyMix (6.25 ml, Roche, catalog number 7958935001) and purified in 1.8 x volume of Agencourt AMPure beads (Beckman Coulter, catalog number A63881). Captured DNA was quantified using QUBIT 2.0. Two wheat captures were pooled (48 libraries) and sequenced in one lane of Illumina NovaSeq S4 (PE150) at the Genome Center of UC Davis.

### Data processing

Illumina sequencing reads were preprocessed to trim adapters with Trimmomatic v0.39 (88). Trimmed reads were aligned to Chinese Spring RefSeqv1.0 (chromosomes A, B and Un) and Kronos De novo assembly using ‘bwa aln’ v0.7.16a-r1181 with a maximum difference 3 (89). Alignments were sorted by using samtools v1.7 (90), and duplicate reads were removed with Picard tools v2.7.1 (http://broadinstitute.github.io/picard/). The promoter capture assay covers two kb of sequence upstream of the start of the annotation of all high-confidence genes of CS RefSeq v1.1 (42). Mutations were called with the MAPS pipeline (*SI Appendix*, Method S2), and those located in residual genetic heterogeneity (RH) regions were removed using a previously published bioinformatics pipeline (5). Error rate determination was performed as described in *SI Appendix*, Method S3. Heterozygous mutations were converted into homozygous classifications when the proportion of wild-type reads was less than 15 % (5). Methods for determining the mutations predicted effects on gene function are described in *SI Appendix*, Method S4.

### Functional validation of mutations in predicted transcription factors’ binding sites

To find conserved elements within the *VRN-A1* promoter, we identified the orthologous genes in other grass species and performed a multiple sequence alignment with the online tool M-Coffee (https://tcoffee.crg.eu/apps/tcoffee/do:mcoffee). Within the conserved region, we searched for potential transcription factor binding sites at PlantPAN 3.0 Promoter Analysis (http://plantpan.itps.ncku.edu.tw/promoter.php). Finally, we studied the effect of these mutations on *VRN-A1* expression using genome-specific qRT-PCR primers developed in a previous study (91), and on spike development and heading time.

## Acknowledgements

We thank Dr. Chengxia Li for her valuable advice on gene expression experiment design, Dr. D.P. Woods for materials for the LFY-binding site mutation experiments, and Huiqiong Lin for helping collect apexes in the LFY-binding site mutation expression experiment. JD acknowledges support from the Howard Hughes Medical Institute (HHMI Researcher Funding, https://www.hhmi.org/) and from the United States Department of Agriculture, National Institute of Food and Agriculture (https://nifa.usda.gov/) competitive Grant 2022-68013-36439 (WheatCAP). HX and JD acknowledge support from the IAEA TC project (CRP5024).

## Data availability

The datasets generated in this study are available in Zenodo (92). The promoter capture data, the re-analyzed exome capture data and the SNP effects can be accessed through the USDA GrainGenes Genome Browsers by selecting the CS reference genome RefSeq v1.0 (https://wheat.pw.usda.gov/GG3/genome_browser). Seeds for the VRN1 promoter Kronos mutants can be requested from the Germplasm Resources Unit at the John Innes Centre and from Dr. Dubcovsky Laboratory Tilling Distribution https://dubcovskylab.ucdavis.edu/wheat-tilling. Backups of the complete Kronos mutant population have been deposited in CIMMYT (Mexico), Shandong Agricultural University in China, the University of Saskatchewan in Canada, the quarantine repository in Australia, the Cereal Disease Laboratory (MN, USA) and Washington State University (WA, USA).

## Notes

### Competing Interest Statement

The authors have declared no competing interest.

