## Supplemental Methods and Tables for "Sequencing 4.3 million mutations in wheat promoters to understand and modify gene expression"

**This file includes:**

Supporting Text, Methods S1 to S5
Tables S1 to S11
Supporting information References 1-6

### Supporting Information Text

#### Method S1. Library preparation and capture.

For the promoter capture, we re-used DNA sequencing libraries generated in our previous exome capture study (1), which included 1,513 $M_2$ plants from the tetraploid wheat cultivar 'Kronos' mutagenized with Ethyl Methane Sulfonate (EMS). Briefly, genomic DNA libraries were constructed using the High-Throughput Library Preparation kits from KAPA Biosystems, Inc. (Wilmington, MA, USA, catalog number KK8234) with dual-SPRI bead size selection. The libraries were barcoded using NEXTflex-96™ oligos 1-48 (Bioo Scientific, Austin, TX, USA, catalogue number 514105) and amplified by PCR (five cycles). The products were purified with Agencourt AMPure beads (Beckman Coulter, A63881) on the Sciclone G3 platform and were eluted in ultrapure DNase/RNase free distilled water and stored at -80 °C.

To capture the gene promoter regions, we used a previously published assay (2) based on 2-kb regions upstream of 110,790 high-confidence wheat genes annotated in Chinese Spring (CS) RefSeq v1.0. Probes for this assay were ordered from Roche (SeqCap EZ Prime Developer Probes 96 Reaction, catalog number 8247633001). For the captures, we pooled 24 DNA libraries together (125 ng/sample, 3 µg in total) and mixed the DNA with blocking oligos for Illumina adapters and repetitive DNA sequences (developer reagent). The DNA mixture was dried using a speed vacuum centrifuge. Capture hybridization and washes were done according to recommended protocols from Roche SeqCapEZ User Guide 4.2 v7. Briefly, the DNA pellet was dissolved in the buffers provided in the SeqCap EZ Hybridization and Wash Kit (Roche, catalog number 5634253001), denatured at 95 °C for 10 minutes, and hybridized to the SeqCap EZ Prime Developer probes for the wheat promoter regions (2) (Roche, catalog number 8247633001) for 70 h at 48 °C in a thermocycler (lid temperature set to 58 °C). The hybridization reaction was recovered using Dynabeads M-270 streptavidin beads (Invitrogen, 653-06) and washed according to the manufacturer's protocol (Roche, catalog number 5634253001). We pooled captures from 48 sequencing libraries and sequenced them in one lane of Illumina NovaSeq S4 (PE150) at the UC Davis Genome Center.

#### Method S2. Identification of SNPs using MAPS.

As a genome reference, we used the CS RefSeq v1.0 genome (3) with the improved CS RefSeq v1.1 annotation. For simplicity, and since the genome coordinates from v1.0 and v1.1 are the same, we will refer to the reference as CS RefSeq v1.1. We aligned the reads from the mutant plants to CS RefSeq v1.1 and called SNPs using default mpileup parameters and mapping quality higher than 20. We then used the MAPS pipeline to select bases in the reference covered by at least one read at quality higher than 20 in a minimum number of samples (MinLib parameter = 38). The minimum number of reads carrying the mutation (minimum coverage, henceforth, MC) required to call that mutation was established independently for homozygous (HomMC) and heterozygous (HetMC). We used four levels of stringency, designated from the lowest to the highest as HetMC3/HomMC2, HetMC4/HomMC3, HetMC5/HomMC3,

HetMC6/HomMC4. For each library, we used the lowest stringency level that resulted in a %EMS > 98%, and we refer to this method as EMS98%. This threshold was sufficient to reduce the percent of non-EMS transitions (A>G/T>C) below 0.7 % in the worst lines. The adjustment of the stringency level by library resulted in the elimination of more SNPs from the low-quality libraries and the inclusion of more mutations from the good quality libraries relative to the fixed HetMC5/HomMC3 stringency level used in the previous exome capture. The new strategy significantly increased the number of called mutations while maintaining a low estimated error.

**Method S3. Error rate determination**

We used four different levels of stringency to call mutations based on the minimum number of reads showing the mutation in heterozygous and homozygous states. The most stringent combination required a heterozygous mutation to be observed in at least six reads and a homozygous mutation to be observed in at least four reads (henceforth, HetMC6/HomMC4). The most liberal criteria required the heterozygous mutation to be observed in at least three reads and a homozygous mutation to be observed in at least two (henceforth, HetMC3/HomMC2). We also used HetMC4/HomMC3 and HetMC5/HomMC3 as intermediate criteria.

The EMS treatment is expected to produce G to A (G>A) or C to T (C>T) mutations (henceforth EMS mutations). At each stringency level, we determined the percent of C to T and G to A mutations, which is designated hereafter as %-EMS. Although we eliminate all the non-EMS mutations from the final database, we cannot eliminate the G>A or C>T errors included among the real mutations induced by the EMS treatment. To estimate the number of these residual errors we used the frequency of reciprocal non-EMS transitions (A>G or T>C SNPs, henceforth, A>G/T>C), which have a similar frequency to the G>A/C>T changes among random mutations (5).

To select a %-EMS threshold, we tested different %-EMS stringency levels and found that ≥ 98% EMS was sufficient to reduce the percent of non-EMS transitions below 0.7 % in the worst lines. We then selected mutations in each line at the first HetMC/HomMC level that had ≥ 98 % EMS mutations. Most of the libraries (940) exceeded the 98%-EMS threshold at the lowest stringency level (HetMC3/HomMC2). Among the other libraries, 389 exceeded the threshold at HetMC4/HomMC3, 120 at HetMC5/HomMC3 and 64 at the highest stringency level (HetMC6/HomMC4). Twenty-two libraries did not reach the 98% EMS threshold even at the highest stringency level (HetMC6/HomMC4) and were eliminated.

**Method S4. Variant Effect Prediction.**

After creating a VCF file with all the EMS mutations, we predicted the effects and SIFT scores (4) of all mutations using the Variant Effect Predictor (VEP) program (5) from Ensembl tools release 103 in offline mode. Release 50 of the *T. aestivum* VEP cache file was downloaded from Ensembl (ftp://ftp.ensemblgenomes.org/pub/plants/release-

50/variation/vep/triticum_aestivum_vep_50_IWGSC.tar.gz). Commands used to install VEP, run the prediction and count the number of different types of mutations were included in the script file 'variant_effects_prediction_for_exome_capture_SNPs.sh' deposited in Zenodo together with the VCF files (doi.org/10.5281/zenodo.7754136).

**Method S5. Estimation of the proportion of mutated "EMS-accessible" G residues.**

The large number of mutations detected in promoter and exome captures provided a unique opportunity to estimate the saturation of our mutant libraries, which equals the proportion of "accessible" GC nucleotides carrying at least one mutation in the population (henceforth, mutated sites). The number of mutations per site follows an approximate Poisson distribution, with most of the sites lacking mutations (n0), a large number carrying one mutation (n1) and rapidly decreasing numbers of sites carrying more than one mutation (n2 to n>7). From our data we can count the number of sites that have one or more mutations, but we need to infer n0 from the Poisson distribution. The main parameter of Poisson distribution is the mean (λ = average number of mutations per site). We estimated the value of λ that maximized the adjustment of the predicted number of sites with mutations in one, two or three lines to the observed values. To do that we selected the λ that minimized the normalized differences between the observed and expected data or $(O-E)^2/E$. In this optimization we excluded the sites with mutation in 4 or more lines to minimize the risk of including residual heterogeneity sites since we observed excess mutation relative to the expected from a Poisson distribution.

Once we optimized λ, we used it to estimate n0 and the accessible GC sites as follows:

λ = total EMS mutations / accessible sites

λ = total EMS mutations / (n0 + mutated sites)

λ = (1 x n1 + 2 x n2 + …) / (n0 + (n1 + n2 + …))

so n0 = (total EMS mutations / λ) - (mutated sites)

and EMS-accessible GC sites = n0 + (mutated sites)

and % saturation of the EMS-accessible space = mutated sites / EMS-accessible GC sites.

### Supporting Tables

**Table S1.** Summary of mutations and uniquely mapped EMS mutations at different stringency levels.

| | Global error | | | | Lib. adjusted |
|---|---|---|---|---|---|
| Stringency | HetMC3/ HomMC2 | HetMC4/ HomMC3 | HetMC5/ HomMC3 | HetMC6/ HomMC4 | EMS 98% |
| Total SNPs | 4,657,354 | 4,120,688 | 3,784,020 | 3,391,163 | 4,345,625 |
| Total EMS mutations * | 4,509,446 | 4,064,952 | 3,749,732 | 3,364,268 | 4,287,361 |
| EMS / kb (all population) | 44.0 | 39.7 | 36.6 | 32.9 | 41.9 |
| EMS / Mb per line | 28.7 | 25.9 | 23.9 | 21.4 | 27.3 |
| Average %-EMS | 96.8 | 98.6 | 99.1 | 99.2 | 98.7 |
| Lowest %-EMS | 34.8 | 72.2 | 91.5 | 95.8 | 98.0 |
| Highest % EMS | 99.49 | 100 | 100 | 100 | 100 |
| Non-EMS transitions † | 11,069 | 7,106 | 5,647 | 4,825 | 9,070 |
| % Estimated error | 0.24 | 0.17 | 0.15 | 0.14 | 0.21% |
| Eliminated RH SNPs | 82,427 | 43,326 | 40,154 | 37,599 | 40,823 |
| % RH | 1.74% | 1.04% | 1.05% | 1.10% | 0.93% |
| Non- mutagenized Kronos | 102 | 46 | 29 | 27 | 0 |
| % EMS | 39.2 | 60.9 | 82.8 | 85.2 | na |

* EMS-type mutations = G>A and C>T, excluding residual heterogeneity (RH).

† Reciprocal transitions A>G and T>C were used to estimate the error rate in EMS mutations.

**Table S2.** Correlations between promoter and exome capture for number of mutations and percent EMS and reciprocal non-EMS transitions across 1,492 common mutant lines at HetMC5/HomMC3 and 1,465 common mutant lines at EMS98%. Correlations were calculated using values obtained with both the previous global threshold HetMC5/HomMC3 for the exome capture remapped to CS RefSeq v1.1 and the new threshold method adjusted by library (EMS98%).

| Trait | HetMC5/HomMC3<br>*R* (n = 1,492) | *P* value | EMS98%<br>*R* (n = 1,465) | *P* value |
|---|---|---|---|---|
| No. EMS | 0.854 | *** | 0.881 | *** |
| %EMS (G>A/C>T) | 0.154 | *** | 0.034 | ns |
| % Reciprocal A>G/T>C | 0.183 | *** | 0.119 | *** |

**Table S3.** Comparison of SNP effects between previous (1) and new exome capture results. The new results are based on mapping the reads to the CS RefSeq v1.1 reference and using the EMS98% stringency method to call mutations. Annotation of SNP effects was performed using VEP from ENSEMBL (Method S3).

| | Krasileva et al. (2017) | New |
|---|---|---|
| Number of Unique EMS mutations * | 4,152,707 | 4,690,454 |
| % EMS | 99.10% | 98.80% |
| Non-EMS-type transitions (estimated error) † | 0.18% | 0.34% |
| Missense variant | 1,030,287 | 1,637,961 |
| Splice acceptor variant | 14,624 | 20,553 |
| Splice donor variant | 15,074 | 20,322 |
| Stop gained | 46,580 | 73,005 |
| Synonymous variant | 550,556 | 921,101 |
| Gene models with at least one mutation (GM1) | 48,172 | 57,950 |
| GM1 with at least one truncation (splice and stop gain) | 28,604 | 38,592 |
| GM1 with at least one missense mutation | 46,198 | 56,792 |
| Avg. number of missense mutations per GM1 | 21.4 | 28.3 |
| GM1 with truncation and/or deleterious missense ‡ | 43,787 | 55,303 |

* EMS-type mutations = G>A and C>T SNPs, excluding residual heterogeneity SNPs

† A>G and T>C SNPs

‡ A SIFT score < 0.05 was used to classify predicted deleterious missense mutations.

**Table S4.** Shared SNPs between promoter capture and exome capture.

| SNP characteristics | Number of lines | Total SNPs |
|---|---|---|
| Promoter Capture | 1,513 | 4,345,625 |
| Exome Capture | 1,521 | 4,748,394 |
| Shared | 1,465 | 174,658 |
| % of Promoter capture | 96.8% | 4.0% |
| % of Exome capture | 96.3% | 3.7% |

**Table S5.** Number of genes and EMS-type SNPs in the promoter and re-mapped exome capture per chromosome.

| **Chromosome** | **HC Gene No. CS RefSeq. v1.1** | **Promoter captures** | **New exome capture** |
|---|---|---|---|
| 1A | 4,344 | 212,380 | 284,009 |
| 1B | 4,714 | 220,188 | 303,578 |
| 2A | 5,797 | 298,433 | 367,898 |
| 2B | 6,109 | 313,699 | 403,738 |
| 3A | 5,235 | 292,665 | 340,485 |
| 3B | 5,905 | 294,564 | 372,226 |
| 4A | 4,861 | 266,762 | 331,289 |
| 4B | 3,869 | 219,342 | 273,502 |
| 5A | 5,429 | 304,837 | 359,177 |
| 5B | 5,576 | 326,000 | 388,667 |
| 6A | 4,127 | 224,092 | 265,065 |
| 6B | 4,615 | 231,492 | 295,899 |
| 7A | 5,552 | 280,522 | 340,698 |
| 7B | 4,855 | 257,069 | 309,623 |
| chrUn | 2,691 | 31,807 | 54,600 |
| Kronos contigs * | Not available | 513,509 | Not available |
| **Total** | 73,679 | 4,287,361 | 4,690,454 |

* The Kronos contigs were used only in the Promoter capture to identify SNPs and were also considered in the sequencing space.

| Correlation | HC Gene No. CS RefSeq. v1.1 | Promoter capture |
|---|---|---|
| Promoter captures | 0.926 | - |
| New exome capture | 0.942 | 0.986 |

**Table S6.** Estimated average percent of GC sites accessible to EMS mutagenesis and proportion of these sites with at least one EMS-type mutation (saturation), using a Poisson approximation.

| Trait | Promoter capture | Exome capture |
|---|---|---|
| Capture sequencing space | 102,378,005 | 127,794,407 |
| % GC in capture sequencing space | 46.8% | 47.3% |
| Total GC sites | 47,861,717 | 60,446,755 |
| Total EMS mutations | 4,287,361 | 4,690,454 |
| Mutated sites | 3,914,089 | 4,343,084 |
| λ that optimizes Poisson distribution | 0.182 | 0.153 |
| Mut. sites in 1 mutant line (Obs. / Exp.) | 3,585,368 / 3,573,950 | 4,032,063 / 4,025,018 |
| Mut. sites in 2 mutant lines (Obs. / Exp.) [*] | 291,577 / 325,229 | 281,686 / 307,914 |
| Mut. sites in 3 mutant lines (Obs. / Exp.) | 31,539 / 19,731 | 25,214 / 15,704 |
| Mut. sites in 4 mutant lines (Obs. / Exp.) | 4,503 / 898 | 3,228 / 601 |
| Estimated No. of GC sites without mutations | 19,642,840 | 26,313,478 |
| Estimated EMS-accessible GC sites | 23,556,929 | 30,656,562 |
| Estimated % EMS-accessible GC sites | 49.2% | 50.7% |
| Estimated % saturation EMS-accessible GC sites [†] | 16.6% | 14.2% |
| % duplicated mutations / total EMS mutations [‡] | 16.4% | 14.0% |

[*] The reduced number of observed mutations relative to the expected from Poisson distribution is likely affected by the MAPS pipeline, which eliminates mutations present in >1 line within the 48 lines included in one sequencing batch to eliminate potential homoeologous SNPs.

[†] Mutated sites / Estimated EMS-accessible GC sites.

[‡] The % saturation estimated from the Poisson adjustment was validated by dividing the number of duplicated mutations present in >1 line by the total number of EMS mutations, which provides a similar probability.

**Table S7.** Effect of a mutation in a conserved SPL binding site on *VRN-A1* expression during spike development. The mutation in the promoter of the *VRN-A1* gene was identified in mutant line K4679. Expression was evaluated in young spikes (5 mm long) in sister lines with and without the mutation. Transcript levels were estimated by qRT-PCR relative to *ACTIN,* which was used as internal control. *VRN-A1* specific qRT-PCR primers were from a previously published study (6). Transcript levels are expressed as fold-*ACTIN* using the ΔCt method.

| | Exp. 1 ($BC_1F_2$) | | Exp. 2 ($BC_1F_3$) | |
|---|---|---|---|---|
| Sample | *SPL*-binding site mutant | WT sister line | *SPL*-binding site mutant | WT sister line |
| 1 | 8.00 | 8.41 | 12.71 | 6.64 |
| 2 | 9.29 | 7.62 | 12.25 | 7.65 |
| 3 | 9.72 | 6.28 | 10.87 | 12.13 |
| 4 | 11.19 | 6.86 | 14.92 | 9.82 |
| 5 | 10.27 | 8.60 | . | . |
| Mean | 9.70 | 7.55 | 12.69 | 9.06 |
| N | 5 | 5 | 4 | 4 |
| S.E.M | 0.53 | 0.44 | 0.84 | 1.22 |
| *t*-test | 0.0145 | | 0.0501 | |

ANOVA combining the two experiments and using experiment as block

***VRN1* transcript levels** relative to *ACTIN* (ΔCt method)

| Source | DF | Sum of Squares | Mean Square | F Value | Pr > F |
|---|---|---|---|---|---|
| Model | 2 | 57.8030 | 28.9015 | 11.31 | 0.0010 |
| Error | 15 | 38.3240 | 2.5549 | | |
| Corrected Total | 17 | 96.1270 | | | |

R-Square: 0.601319

| Source | DF | Type III SS | Mean Square | F Value | Pr > F |
|---|---|---|---|---|---|
| Exp | 1 | 22.4950 | 22.4950 | 8.80 | 0.0096 |
| Genotype | 1 | 35.3080 | 35.3080 | 13.82 | **0.0021** |

**Normality of residuals:** Shapiro-Wilk test Pr < W = 0.9124

Least Squares Means

| Genotype | qRT-PCR LSMEAN | Standard Error |
|---|---|---|
| WT | 8.348 | 0.534 |
| mut | 11.149 | 0.534 |

*VRN-A1* transcript levels in the SPL-binding site mutant were 33.6 % higher than in the wildtype sister line.

**Table S8.** Phenotypic effect of a mutation within a conserved SPL-binding site within the *VRN-A1* promoter. The mutation was identified in line K4679, and its effect on days to heading (DTH) and spikelet number per spike (SNS, avg. two taller tillers) was evaluated in two experiments.

| | Exp. 1 DTH | | Exp. 2 DTH | | Exp. 1 SNS | | Exp. 2 SNS | |
|---|---|---|---|---|---|---|---|---|
| Sample | *SPL*-bind. site mut. | WT sister line | SPL-bind. site mut. | WT sister line | SPL-bind. site mut. | WT sister line | SPL-bind. site mut. | WT sister line |
| 1 | 46 | 45 | 48 | 51 | 16.0 | 16.0 | 16.0 | 15.5 |
| 2 | 45 | 45 | 53 | 49 | 15.5 | 14.0 | 15.0 | 15.0 |
| 3 | 46 | 44 | 46 | 52 | 16.0 | 15.5 | 14.5 | 16.0 |
| 4 | 44 | 44 | 52 | 50 | 15.0 | 13.5 | 16.0 | 15.0 |
| 5 | 45 | 45 | 50 | 50 | 16.0 | 15.5 | 16.0 | 16.0 |
| 6 | . | . | 49 | 48 | . | . | 15.0 | 14.5 |
| 7 | . | . | 49 | 46 | . | . | 16.0 | 14.0 |
| 8 | . | . | 46 | 48 | . | . | 14.0 | 15.5 |
| Mean | 45.20 | 44.60 | 49.13 | 49.25 | 15.70 | 14.90 | 15.31 | 15.19 |
| S.E.M | 0.37 | 0.24 | 0.90 | 0.67 | 0.20 | 0.48 | 0.28 | 0.25 |
| *t*-tests | 0.2165 | | 0.9128 | | 0.1656 | | 0.7448 | |

ANOVAs combining the two experiments and using experiment as block

**Days to heading**

| Source | DF | S. Squares | Mean Square | F Value | Pr > F |
|---|---|---|---|---|---|
| Model | 2 | 113.27788 | 56.63894 | 17.33 | <.0001 |
| Error | 23 | 75.18365 | 3.26885 | | |
| Corrected Total | 25 | 188.46154 | | | |

R-Square: 0.601066

| Source | DF | Type III SS | Mean Square | F Value | Pr > F |
|---|---|---|---|---|---|
| Exp | 1 | 113.12404 | 113.12404 | 34.61 | <.0001 |
| Genotype | 1 | 0.15385 | 0.15385 | 0.05 | **0.8302** |

**Normality of residuals:** Shapiro-Wilk test: Pr < W = 0.3098

Least Squares Means

| Genotype | DTH LSMEAN | Standard Error |
|---|---|---|
| WT | 46.96683 | 0.50845 |
| mut | 47.12067 | 0.50845 |

**Spikelet number per spike** (power transformation **10 to restore Normality)

| Source | DF | Sum of Squares | Mean Square | F Value | Pr > F |
|---|---|---|---|---|---|
| Model | 2 | 1.78E23 | 8.903E22 | 0.86 | 0.4350 |
| Error | 23 | 2.378E24 | 1.031E23 | | |
| Corrected Total | 25 | 2.554E24 | | | |

R-Square: 0.069828

| Source | DF | Type III SS | Mean Square | F Value | Pr > F |
|---|---|---|---|---|---|
| Exp | 1 | 1.116E22 | 1.116E22 | 0.11 | 0.7452 |
| Genotype | 1 | 1.669E23 | 1.669E23 | 1.62 | **0.2160** |

**Normality of residuals:** Shapiro-Wilk test: Pr < W = 0.0511

Least Squares Means (original scale not transformed)

| Genotype | SNS LSMEAN | Standard Error |
|---|---|---|
| WT | 15.0826923 | 0.2204898 |
| mut | 15.4673077 | 0.2204898 |

Mutations in the putative SPL binding site were not associated with significant differences in DTH and SNS

**Table S9.** Effect of a mutation in a conserved LFY-binding site on *VRN-A1* expression during spike development (Waddington 3.25). The induced EMS mutation was identified in line K2944 in the *VRN-A1* promoter. Transcript levels were estimated by qRT-PCR relative to *ACTIN,* which was used as internal control. *VRN-A1* specific qRT-PCR primers were from a previously published study (6). Transcript levels are expressed as fold-*ACTIN* using the ΔCt method.

| | Exp. 1 *VRN-A1* expression | | Exp. 2 *VRN-A1* expression | |
|---|---|---|---|---|
| Sample | *LFY*-binding site mut. | WT sister line | *LFY*-binding site mut. | WT sister line |
| 1 | 1.52 | 3.29 | 6.83 | 5.13 |
| 2 | 3.73 | 1.36 | 5.72 | 4.56 |
| 3 | 3.86 | 1.39 | 5.57 | 5.81 |
| 4 | 5.72 | 2.36 | 5.48 | 5.39 |
| 5 | . | . | 6.11 | 5.00 |
| 6 | . | . | 4.99 | 5.48 |
| 7 | . | . | 5.67 | 5.12 |
| 8 | . | . | 6.41 | 4.70 |
| Mean | 3.71 | 2.10 | 5.85 | 5.15 |
| S.E.M | 0.86 | 0.46 | 0.20 | 0.14 |
| *t*-test | 0.1499 | | 0.0143 | |

***VRN1* transcript levels** relative to *ACTIN* (ΔCt method). Experiment was used as block.

| Source | DF | Sum of Squares | Mean Square | F Value | Pr > F |
|---|---|---|---|---|---|
| Model | 2 | 41.9175 | 20.9588 | 27.48 | <.0001 |
| Error | 21 | 16.0178 | 0.7628 | | |
| Corrected Total | 23 | 57.9353 | | | |

R-Square: 0.6312

| Source | DF | Type III SS | Mean Square | F Value | Pr > F |
|---|---|---|---|---|---|
| Exp | 1 | 35.8975 | 35.8975 | 47.06 | <.0001 |
| Genotype | 1 | 6.0200 | 6.0200 | 7.89 | 0.0105 |

**Normality of residuals.** Shapiro-Wilk test Pr < W = 0.3612

Least Squares Means

| Genotype | Vrn1 LSMEAN | Standard Error |
|---|---|---|
| WT | 3.7001 | 0.2599 |
| M | 4.7018 | 0.2599 |

*VRN-A1* transcript levels in the LFY-binding site mutant are on average 27.1% higher than in the wildtype sister line.

**Table S10.** Phenotypic effects of the K2944 mutation in a conserved LFY-binding site within the *VRN-A1* promoter. The effect on spikelet number per spike (SNS) in the main shoot was evaluated in two independent experiments using $BC_1F_2$ and $BC_1F_3$ plants (derived from a single heterozygous $BC_1F_2$ plant). The effect on days to heading (DTH) and leaf number (LN) was evaluated in one experiment using $BC_1F_3$ plants.

#### Spikelet number per spike

| | Exp. 1. Spikelet No. / spike | | | Exp. 2. Spikelet No. / spike | |
|---|---|---|---|---|---|
| | mutant | WT | | mutant | WT |
| | 13 | 15 | 14 | 15 | 16 |
| | 14 | 14 | 14 | 15 | 16 |
| | 12 | 14 | 15 | 15 | 15 |
| | 14 | 14 | 14 | 16 | 16 |
| | 15 | 15 | | 14 | 17 |
| | 12 | 14 | | 17 | 17 |
| | 13 | 14 | | 14 | 17 |
| | 13 | 14 | | | 16 |
| | 12 | 14 | | | 17 |
| | 14 | 14 | | | 17 |
| | | 13 | | | 16 |
| | | 14 | | | 15 |
| Average | 13.2 | 14.1 | | 15.1 | 16.3 |
| n | 10 | 16 | | 7 | 12 |
| S.E.M | 0.33 | 0.13 | | 0.40 | 0.22 |
| *t*-test | 0.0046 | | | 0.0168 | |
| Kruskal-Wallis [1] | 0.0113 | | | not necessary | |

[1] In Exp 1 no transformation restored normality of residuals so we used a non-parametric test

**Average spikelet number per spike (SNS)**

```
Data transformed using a power transformation (**-2) to improve normality of residuals.
For the untransformed data the P values for Genotype was 0.0002.

                                      Sum of
Source                       DF      Squares      Mean Square     F Value    Pr > F
Model                         2    0.0000242       0.0000121       39.67     <.0001
Error                        42    0.0000128       0.0000031
Corrected Total              44    0.0000370
R-Square: 0.65386

Source                       DF   Type III SS     Mean Square     F Value    Pr > F
Experiment                    1    0.0000184       0.0000184       59.00     <.0001
Genotype                      1    0.0000055       0.0000055       17.99     0.0001

Normality of residuals. Shapiro-Wilk test Pr < W      0.0129

Least Squares Means (untransformed)
                           Standard
Genotype      SNS LSMEAN      Error
M              14.1815       0.1940 6.6% decrease
WT             15.1826       0.1512
```

### Leaf number and days to heading

| | Exp. 2. Leaf Number | | Exp. 2. Days to heading | |
|---|---|---|---|---|
| | mutant | WT | mutant | WT |
| | 8 | 8 | 46 | 45 |
| | 9 | 9 | 46 | 46 |
| | 9 | 9 | 46 | 46 |
| | 9 | 8 | 47 | 46 |
| | 8 | 9 | 46 | 46 |
| | 9 | 9 | 47 | 46 |
| | 9 | 10 | 46 | 49 |
| | | 9 | | 47 |
| | | 9 | | 46 |
| | | 9 | | 47 |
| | | 8 | | 46 |
| | | 9 | | 45 |
| Average | 8.7 | 8.8 | 46.3 | 46.3 |
| n | 7 | 12 | 7 | 12 |
| S.E.M | 0.18 | 0.16 | 0.18 | 0.30 |
| *t*-test | 0.6533 | | 0.9343 | |
| Kruskal-Wallis [1] | 0.6780 | | 0.5888 | |

The ANOVAs using the parametric analyses was not significant for both leaf number and days to heading (*t*-Test above) but the residuals were not normally distributed, so we performed non-parametric Kruskal-Wallis test, which were also not significant:

#### Leaf number

```
Kruskal-Wallis Test
Chi-Square            0.1723
DF                         1
Pr > Chi-Square       0.6780
```

#### Days to heading

```
Kruskal-Wallis Test
Chi-Square            0.2922
DF                         1
Pr > Chi-Square       0.5888
```

**Table S11.** Relative abundance and % of heterozygous mutations among non-EMS errors.

| | Exome Capture | | | | Promoter Capture | | | |
|---|---|---|---|---|---|---|---|---|
| Type | Het | Hom | % Het | % of errors | Het | Hom | % Het | % of errors |
| **EMS mutations** | | | | | | | | |
| CT | 1,530,272 | 816,553 | 65.21 | | 1,390,714 | 756,335 | 64.77 | |
| GA | 1,521,660 | 821,969 | 64.93 | | 1,382,114 | 758,198 | 64.58 | |
| % het. EMS mut. | 3,051,932 | 1,638,522 | 65.07 | | 2,772,828 | 1,514,533 | 64.67 | |
| **Non-EMS errors** | | | | | | | | |
| AC | 2,097 | 0 | 100.0 | 3.6 | 4,070 | 1 | 100.0 | 7.0 |
| TG | 2,026 | 0 | 100.0 | 3.5 | 3,282 | 0 | 100.0 | 5.6 |
| AG | 7,756 | 1 | 100.0 | 13.4 | 4,539 | 0 | 100.0 | 7.8 |
| TC | 7,847 | 1 | 100.0 | 13.5 | 4,528 | 3 | 99.9 | 7.8 |
| AT | 5,849 | 0 | 100.0 | 10.1 | 5,528 | 0 | 100.0 | 9.5 |
| TA | 5,937 | 0 | 100.0 | 10.2 | 5,457 | 1 | 100.0 | 9.4 |
| CG | 2,518 | 0 | 100.0 | 4.3 | 1,468 | 1 | 99.9 | 2.5 |
| GC | 2,591 | 0 | 100.0 | 4.5 | 1,457 | 0 | 100.0 | 2.5 |
| CA | 10,660 | 0 | 100.0 | **18.4** | 13,564 | 1 | 100.0 | **23.3** |
| GT | 10,655 | 2 | 100.0 | **18.4** | 14,364 | 0 | 100.0 | **24.7** |
| % het. non-EMS mut. | 57,936 | 4 | 99.99 | | 58,257 | 7 | 99.99 | |
